# LAG3 is a Central Regulator of NK Cell Cytokine Production

**DOI:** 10.1101/2020.01.31.928200

**Authors:** Sriram Narayanan, Patricia J. Ahl, Veonice Au Bijin, Nivashini Kaliaperumal, Seng Gee Lim, Cheng-I Wang, Anna-Marie Fairhurst, John E. Connolly

## Abstract

Natural killer (NK) cells are innate effectors, which play a crucial role in controlling viral infections. Administration of IFN-α has shown promising results as a therapeutic, controlling HIV, and chronic viral hepatitis. However the downstream mechanisms by which IFN-α mediates its anti-viral effects is largely unknown. In this investigation, we evaluated the impact of IFN-α on peripheral blood NK cells from healthy donors. High dimensional flow cytometry analysis of NK cell surface receptors following exposure to IFN-α showed an increased expression of the check point inhibitor LAG3. Further characterization revealed that LAG3 was expressed in a subset of NK cells with high expression of activation and maturation markers. Assessment of metabolic pathways showed that LAG3+ NK cells had enhanced rates of glycolysis and glycolytic capacity, suggesting that it is a primed effector subset with enhanced glucose metabolism. Inhibition of LAG3 on NK cells using antibody *in vitro* resulted in a profound increase in secretion of cytokines IFN-γ, TNF-α, MIP-1α and MIP-1β, without affecting the cytotoxic activity. Taken together, these results showed that LAG3 is a negative regulator of cytokine production by mature NK cells.

## Introduction

The plasmacytoid (pDC)-Natural Killer (NK) cell axis acts as a first line of defence by the host against viral infections. pDCs secrete type I interferons upon recognizing virus-associated pathogen associated molecular patterns (PAMPs) through pattern recognition receptors (PRRs). Type I interferons (IFN-I) bind to IFN (IFN-α/β) receptors on the surface of NK cells to prime, activate and initiate mechanisms for the destruction of infected cells. The subsequent stimulation of the IFN signalling pathway results in the increased expression of several interferon stimulated genes (ISGs) which have direct and indirect antiviral activities (1).

In chronic viral infections, such as hepatitis B, or C virus (HBV or HCV respectively), NK cells demonstrate reduced cytotoxicity and cytokine secretion (2,3) Additionally, a reduction in circulating numbers of NK cells has been observed (2), together with reduced levels of their activating receptors (4). Type I interferons produced during a viral infection and cytokines, such as IL-12, IL-15 and IL-18, prime NK cells to enhance their cytotoxic and cytokine secretory functions (5). Therapeutic administration of PEGylated-IFN-α results in loss of HBsAg, a hallmark of functional cure in chronic HBV patients (6). IFN-α therapy resulted in increased surface expression of activating receptors on NK cells, including NKp46, TRAIL. NK cells also showed restored cytotoxicity and production of IFNγ (7–9). Taken together, these studies strongly support a direct role for NK cells in controlling viral infections mediated by IFN-α stimulation.

NK cells express a wide variety of activating and inhibitory receptors and their function is determined by the balance between positive and negative signaling from these receptors. It is crucial, therefore, to understand the phenotype of different NK cell subsets during disease and treatment. We aimed to directly ascertain the phenotypic and functional changes induced in NK cells following IFN-α stimulation. Exposure to IFN-α induced the expression of LAG3, an inhibitory receptor, commonly associated with T cell exhaustion in chronic infections and cancer. To date, the function of LAG3 in NK cells is not well understood due to contrasting results from mouse and human NK cells. While studies using knockout mouse models suggested a positive role for LAG3 in NK cell cytotoxicity, these could not be successfully reproduced in human NK cells (10,11). High dimensional flow cytometry analysis in these studies revealed that LAG3 expression marks a subset of mature activated NK cells. Metabolic assays additionally showed that NK cells expressing LAG3 have enhanced glycolytic activity. Blockade of LAG3 enhanced cytokine production by activated NK cells without altering their cytotoxicity. Taken together, these results demonstrate that LAG3 is a negative regulator of cytokine production and a potential therapeutic target for chronic viral infections.

## Materials and methods

### Isolation and stimulation of NK cells

Blood samples from healthy volunteer donors were obtained after approval by the National Healthcare Group Domain Specific Review Board, Singapore (NHG DSRB Ref: 2000/00828). Samples were collected after getting written consents from volunteers. PBMCs were isolated from blood by Ficoll density gradient centrifugation (Ficoll-Paque, GE Healthcare). NK cells were isolated using the NK cell isolation kit (Miltenyi Biotec) following the manufacturer’s protocol. Purity was 94% to 99%, confirmed by flow cytometry using the markers, CD3 and CD56. Isolated NK cells were stimulated for 16 hours *in vitro* with recombinant IL-2 (100 U/ml) or recombinant IFN-α (10 ng/ml) or a combination of both, in Iscove’s Modified Dulbecco’s Medium (IMDM, Gibco) supplemented with 10% FBS, 100U/ml penicillin, 100 μg/ml streptomycin, 1mM pyruvate, 2mM L-glutamine, nonessential amino acids and 15mM HEPES (complete (cIMDM)). Cells left untreated in medium alone were used as a negative control. For flow cytometry analysis, PBMCs were stimulated for different time periods up to 96 hours, as described in the figure legends.

NK cells were expanded by culturing with IFN-α and IL-2 (Miltenyi Biotec) in cIMDM for 48 hours to induce priming. They were then co-cultured with genetically modified K562 cells, expressing 4-1BBL, membrane bound IL-21 and CD86 (TD reference number GIS/Z/10345) (28), at a ratio of 1:1 (NK:K562) in cIMDM with IL-2 (100 U/ml) and IL-15 (10 ng/ml) for up to 3 weeks. Fresh medium containing IL-2 and IL-15 was added to the cells every 3 days. The cells were then washed, counted, stained and sorted as described below.

### RNA isolation and gene expression analysis

The cells were washed in PBS and the cell pellet was lysed in RLT lysis buffer containing β-Mercapto Ethanol and stored at −80°C. Total RNA was extracted from cell lysates using RNeasy Mini Kit (Qiagen, Valencia, CA, USA), according to the manufacturers’ instructions. Total RNA (50ng/hybridization reaction) was used to analyze the immune gene expression profile in NK cells using the nCounter^®^ Human Immunology v2 Panel (NanoString Technologies, Seattle, WA, USA). This technology relies on the direct hybridization of total RNA with fluorescent barcoded probes complementary to 579 immune response-related genes and 15 internal reference controls. The hybridized probes were purified and counted on the nCounter^®^ SPRINT Profiler (NanoString Technologies). Data was analysed with the nSolver 4.0 software (NanoString Technologies). After subtracting background noise (calculated as the mean plus 2 SD of the negative control probes, excluding outliers), intra-/inter-assay variability was adjusted according to the signal of the positive control probes. The data were normalized against the ratio between the geometric mean of all positive controls and the geometric mean of the positive controls of each sample. The samples were then normalized for RNA content according to the relative expression of the housekeeping genes provided in the assay.

### Antibodies and flow cytometry

The following antibodies were used in NK cell flow cytometry panel; CD56 PE-CF594 (NCAM16.2), CD3 BV570 (UCHT1), CD14 APC-Cy7 (M5E2), CD19 APC-Cy7 (HIB19), CD57 BUV615 (NK-1), LAG3 PE-Cy7 (T47-530), CD226 BUV395 (DX11), CD161 BB790 (DX12), CD16 BUV805 (3G8), CD69 BB660 (FN50), CD94 BB630 (HP-3D9), NKG2D PerCP-Cy5.5 (1D11), CD319 APC (235614), CCR5 BB515 (DX26), NKp30 BUV563 (P30-15), NKp46 BUV661 (9E2/Nkp46), NKp44 BV650 (p44-8), CD158e1 BV780 (DX9), PD-1 BUV737 (42D1), CD253 (TRAIL) BV421 (RIK-2), CXCR6 BV480 (13B1E5), CD158b BV605 (CH-L), TIM3 (CD366) BV750 (7D3), CD38 PE-Cy5 (HIT2) (all from BD Biosciences), NKG2C PE (FAB138P), NKG2A AlexaFluor700 (FAB1059N) (from R&D systems). All antibodies were titrated and used at optimal dilutions prior to panel integration. Single-cell suspensions of PBMCs were first stained for dead cells using LIVE/DEAD™ Fixable Near-IR Dead Cell Stain Kit (Thermo Fisher) in PBS for 10 minutes and then blocked for 15 minutes in FACS staining buffer (phosphate buffered saline with 1% fetal calf serum and 15 m*M* HEPES) containing FcR-blocking antibody (eBioscience). Cells were then incubated on ice for 30 minutes with the mixture of antibodies from the panel, washed and re-suspended in FACS staining buffer. Samples were acquired using a FACSymphony™ cell analyser (Beckton Dickinson). Data were analysed using FlowJo 10 for Windows (Tree Star). A positive value was calculated using a fluorescence minus one (FMO) control for each of the antibodies used. The gating strategy used for NK cell analysis is shown in Supplementary figure 2A. High dimensional analyses were performed using the UMAP dimensionality reduction algorithm plug-in from FlowJo and cell-cluster analysis was carried out using the Phenograph plug-in. Heatmapper web tool (www.heatmapper.ca) was used for construction of heatmaps from Phenograph cell cluster data. Expanded NK cells were sorted based on LAG3 expression using a BD Aria II (Beckton Dickinson) and post sort showed more than 97% purity for the sorted populations.

### NK cell cytotoxicity and cytokine secretion

NK cells were co-cultured with THP1 cells loaded with Calcein-AM dye, at different Effector:Target (E:T) ratios, in a 96 well plate for 5 hours. Cytotoxicity was measured by the fluorescence of Calcein dye released in the culture medium upon lysis of THP1 cells. Target cells cultured in the absence of NK cells was used as negative control. Cytokines secreted in the supernatant was measured using Luminex. For LAG3 blocking experiments, anti-LAG3 antibody or control antibody (Dr. Cheng I-Wang, SIgN, A*STAR) was added to NK cells (10 ug/ml) and target cells and proceeded with assay as above.

### Metabolic analysis of NK cells

NK cells were expanded *in vitro* and sorted based on LAG3 expression were stimulated in complete RPMI and plated at 200,000 cells per well. Glycolytic function and mitochondrial respiration were measured by extracellular acidification rate (ECAR, mpH/min) and oxygen consumption rate (OCR, pmol/min) respectively, using XFe96 extracellular flux analyzer (Seahorse Bioscience). Respiration was measured in XF Assay Modified Media with L-glutamine (2mM), Sodium Pyruvate (1mM) and with or without Glucose (11mM) for OCR and ECAR measurements, respectively.

### Statistical analysis

Data were analysed using GraphPad Prism 7.01 for Windows. Error bars in graphs indicate SEM. Nonparametric data were assessed using the Mann-Whitney test for 2 comparisons and the Kruskal-Wallis test with post hoc Dunn’s multiple comparisons test for 3 or more comparisons unless otherwise stated. The significant differences are indicated by *p<0.05, **p<0.01, ***p<0.001.

## Results

### Interferon-α induces LAG3 mRNA expression in NK cells

Little is known about the mechanism behind enhanced NK responses upon priming with IFNα. To unravel this, NK cells were isolated from healthy donor PBMCs and then cultured with IFN-α in the presence or absence of IL-2. The latter was added to maintain the survival of NK cells. After 16 hours, gene expression was analysed using Nanostring. The presence of IFN-α resulted in the induction of a distinct set of genes compared to IL-2 stimulation alone (Fig 1a). As expected, known IFN-α inducible genes, which included *MX1*, *IFIT2*, *IFI35*, *IFITM1* and *IFIH5*, were upregulated in this group (Fig 1A). Importantly, the expression of the NK cell activating receptors, *TRAIL* and *SLAMF7*, were increased in response to IFN-α (Fig 1B and C). Analyses of inhibitory receptors revealed that among all the inhibitory receptors, *LAG3* mRNA was uniquely upregulated among all the other inhibitory receptors measured in response to IFN-α (Fig 1B and C). Expression of *LAG3* mRNA was 15 fold more than the average expression of *TIM3*, the second most upregulated inhibitory receptor, induced by IL-2 and IFN-α. These results showed that *LAG3* gene expression is induced by IFN-α in NK cells.

**Figure 1:**
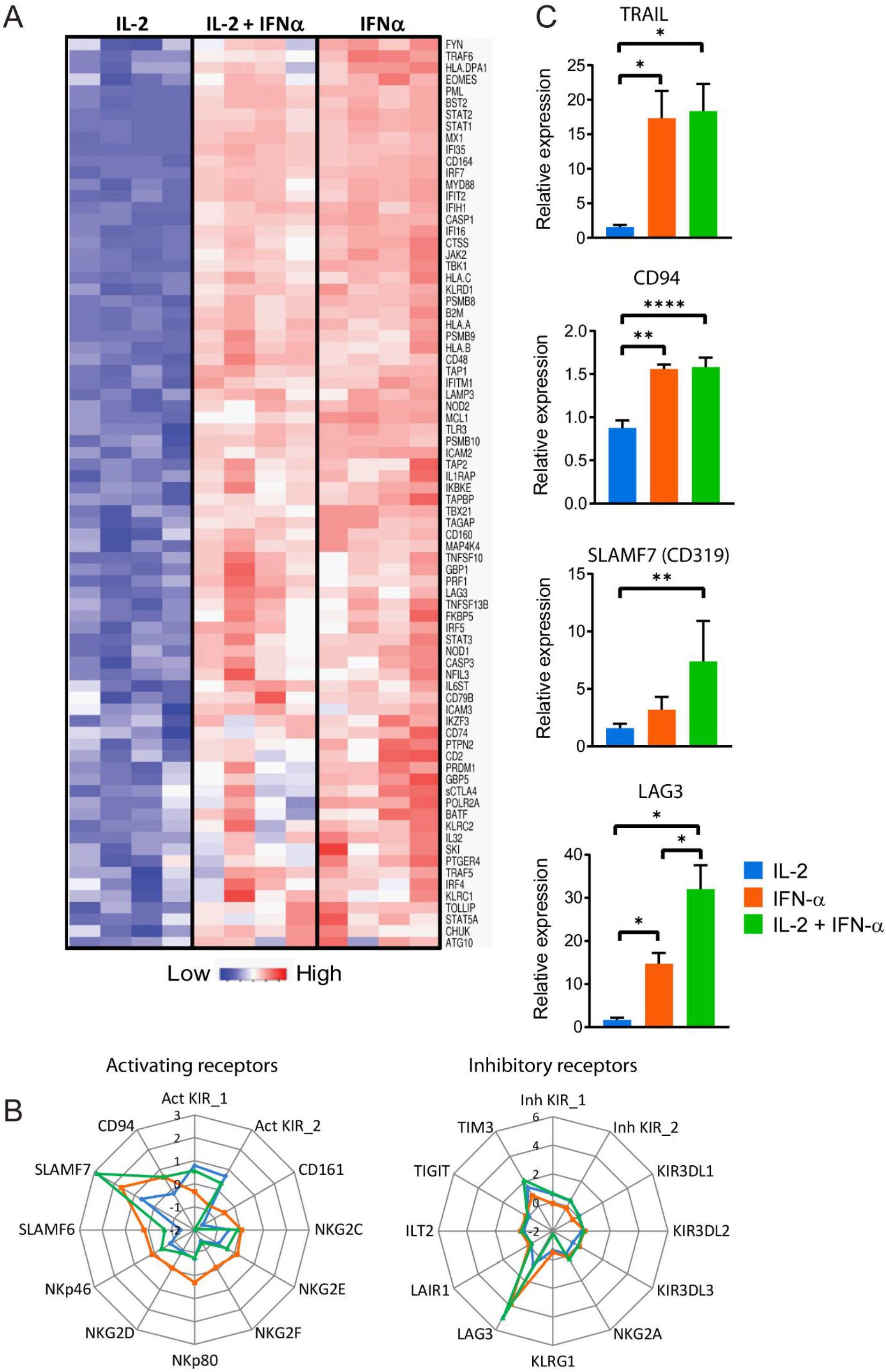
NK cells upregulated LAG3 mRNA in response to IFNα. NK cells were cultured in vitro with IL-2 or IFNα, or IL-2/ IFN-α for 16 hours and mRNA levels analysed by Nanostring^®^. **(A)** Heat map showing the expression of genes induced by IFN-α with or without IL-2 relative to NK cells that were left unstimulated. **(B)** Radar plots showing the relative expression levels (as log 2 values) of various activating receptors (left) and inhibitory receptors (right) expressed in NK cells treated as shown in legend. **(C)** Relative expression of selected genes that are induced by IFNα. Dotted line at 1.0 on y-axis indicates untreated mRNA relative expression. Data shown is from 4 different donors.

### NK cells upregulate surface expression of LAG3 following *IFN-α* exposure

We went on to examine the expression of LAG3 at single cell level. PBMCs from healthy donors were stimulated with IFN-α in the presence or absence of IL-2 for various time intervals and the cell surface expression of LAG3 was analysed by flow cytometry. NK cells were analysed after gating out dead cells, monocytes and B cells as shown in the gating strategy in Supp. figure 2A. An analysis of the kinetics of LAG3 expression showed that the frequency of LAG3+ NK cells increased with IFN-α and IL-2 combination treatment as early as 24 hours and was maintained up to the time point studied (Fig 2A). LAG3 expression on NK cells was induced significantly by IFN-α alone or IFN-α and IL-2, but not by IL-2 alone, when compared to unstimulated cells 48 hours post stimulation (Fig 2B). Protein expression of other receptors like TRAIL, CD319, CD94 and TIM3 was also increased with IFN-α stimulation corroborating gene expression analysis (Supp. figure 2B). Analysis of NK subsets revealed a higher frequency of LAG3 positivity (LAG3+) in the CD56^low^ population compared to CD56^high^ NK cells. However, no difference in LAG3 expression was observed with respect to CD16 expression on NK cells (Fig 2C) suggesting that LAG3 expression is not a general phenomenon of activation with cytokines on all NK cell subsets. Our analysis showed that at single cell level, LAG3 receptor is preferentially expressed on CD56^low^ subset of NK cells.

**Figure 2:**
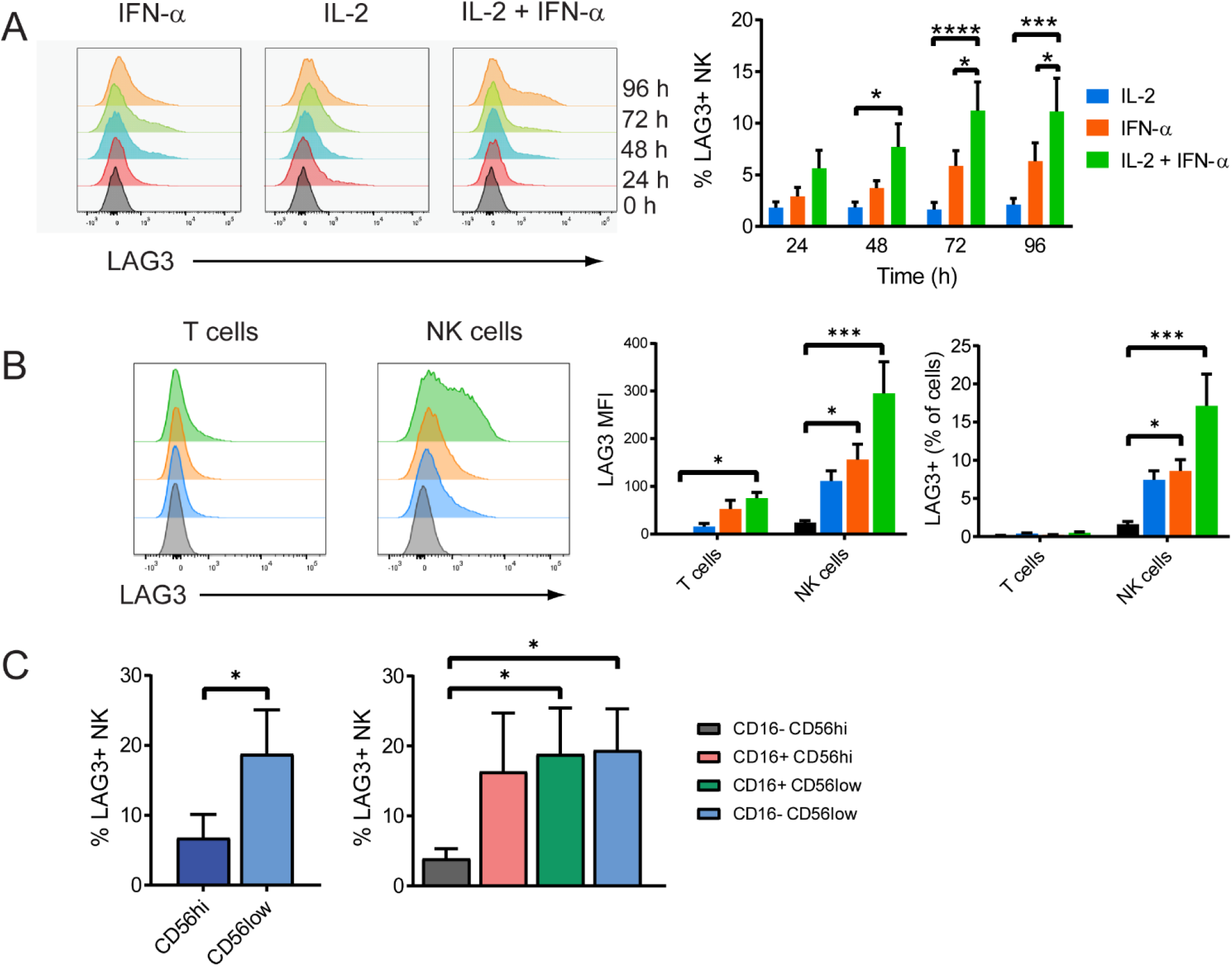
Kinetics of LAG3 receptor expression induced by IFN-α in NK cells. **(A)** PBMCs stimulated under various conditions for different time points indicated were analysed for the expression of LAG3 by flow cytometry. N=5 different donors**. (B)** Cellular expression levels of LAG3 and frequency of LAG3+ cells in NK and T cells from PBMCs stimulated with IFN-α and IL-2 for 48 hours are shown. Representative histograms of LAG3 expression NK cells cultured for 48 hours. **(C)** Expression of LAG3 in different NK cell subsets from cells cultured with IL-2 and IFNα. N=5 different donors.

### LAG3 expression marks a mature and activated NK cell subset

Previous work has shown that LAG3 is expressed on T cells following chronic antigen stimulation (12). Additionally, it has been characterised as a marker of T cell exhaustion, together with PD-1, in HIV and cancer (13,14). Little is known about the ontogeny and function of LAG3 in human NK cells. Therefore, to understand the nature of LAG3+ NK cells we went on to culture PBMCs from healthy donors with IFN-α /IL-2 and then evaluated the expression of activating and inhibitory receptors on this subpopulation. LAG3+ NK cells had higher levels of CD57, NKG2C, CD158b and TIM3, compared to LAG3-NK cells (Fig 3A). Similarly, the frequency of LAG3+ cells was higher in mature CD57+ NK subset (Supp. Figure 3A). The increased expression of CD57 and CD158b suggested that LAG3+ NK cells are more mature than LAG3-NK cells (15) and elevated surface expression of TIM3 indicated that LAG3+ NK cells were activated (16).

**Figure 3:**
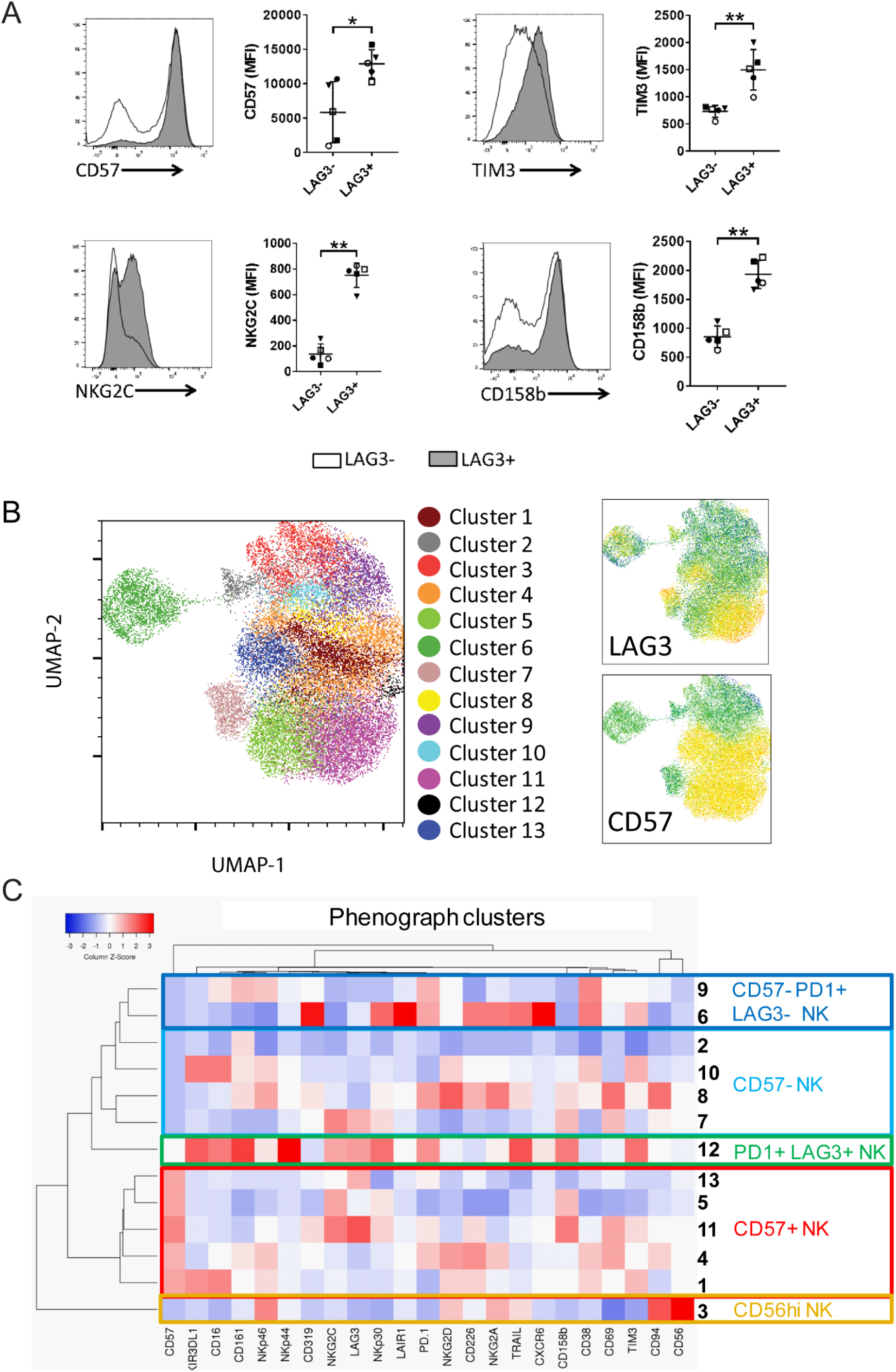
LAG3+ NK cells have higher expression of activation and maturation markers. **(A)** NK cells stimulated with IFN-α for 48 hours were gated for LAG3 expression and analysed for expression of other activating and inhibitory receptors by flow cytometry. Representative histograms depicting expression of markers that differ significantly between LAG3+ and LAG3-NK cells are shown (left panel). Profiles from LAG3- and LAG3+ NK cells are represented by unshaded and shaded histograms respectively. The quantitative expression from multiple donors is shown (right panel). **(B)** UMAP clustering analysis of concatenated NK cells from flow cytometry data. The clusters of NK populations identified by phonograph and their LAG3 and CD57 expression intensities are shown. **(C)** Heat map of the expression of various markers in the clusters are shown. The highlighted clusters indicate NK cell subsets of mature CD57+ phenotype (red), CD57-phenotype (light blue), LAG3+ and LAG3-exhausted phenotype (green and dark blue, respectively) and CD56high NK cells (orange). N=5 different donors.

Further characterization using the UMAP dimensionality reduction algorithm revealed that LAG3 expression was on multiple NK cell subsets (Fig 3B). Heat map analysis of phenograph clusters indicated that LAG3 was expressed predominantly on CD57+ mature NK cells (Fig 3C). These clusters formed the majority of LAG3 expressing subsets, and also co-expressed the activating receptors CD319, NKG2C, CD226 and CD38 and the inhibitory receptor TIM3 (Fig 3C and Supp. Figure 3B). LAG3 expression was also observed on a small population of NK cells expressing lower levels of CD57 and higher levels of PD-1 and TIM3, which are considered to be markers of T cell exhaustion. These results confirmed that LAG3 expression is associated with NK cell activation and maturation.

### LAG3 expression defines a subset of NK cells with increased glycolytic activity

Metabolic reactions play a key role in defining immune cell function. Therefore, we went on to evaluate real-time metabolic flux on sorted LAG3+ and LAG3-NK cell subsets using a Seahorse Analyzer. LAG3+ NK cells exhibited higher glycolytic function compared to LAG3-NK cells (Fig. 4A). Specifically, LAG3 expressing NK cells had higher rate of glycolysis and higher glycolytic capacity than LAG3-NK cells (Fig. 4B-C). Both indicate enhanced utilization of glycolytic pathway in the LAG3+ cells. No significant differences were observed for glycolytic reserve and non-glycolytic acidification between the LAG3- and LAG3+ subsets (Supp. Fig 4). Furthermore the OCR was similar between the two subsets, indicating no change in the oxidative phosphorylation pathway (Supp. Fig 4B). These results indicate that LAG3 expression defines a subset of NK cells that are more glycolytically active compared to LAG3-NK cells.

**Figure 4:**
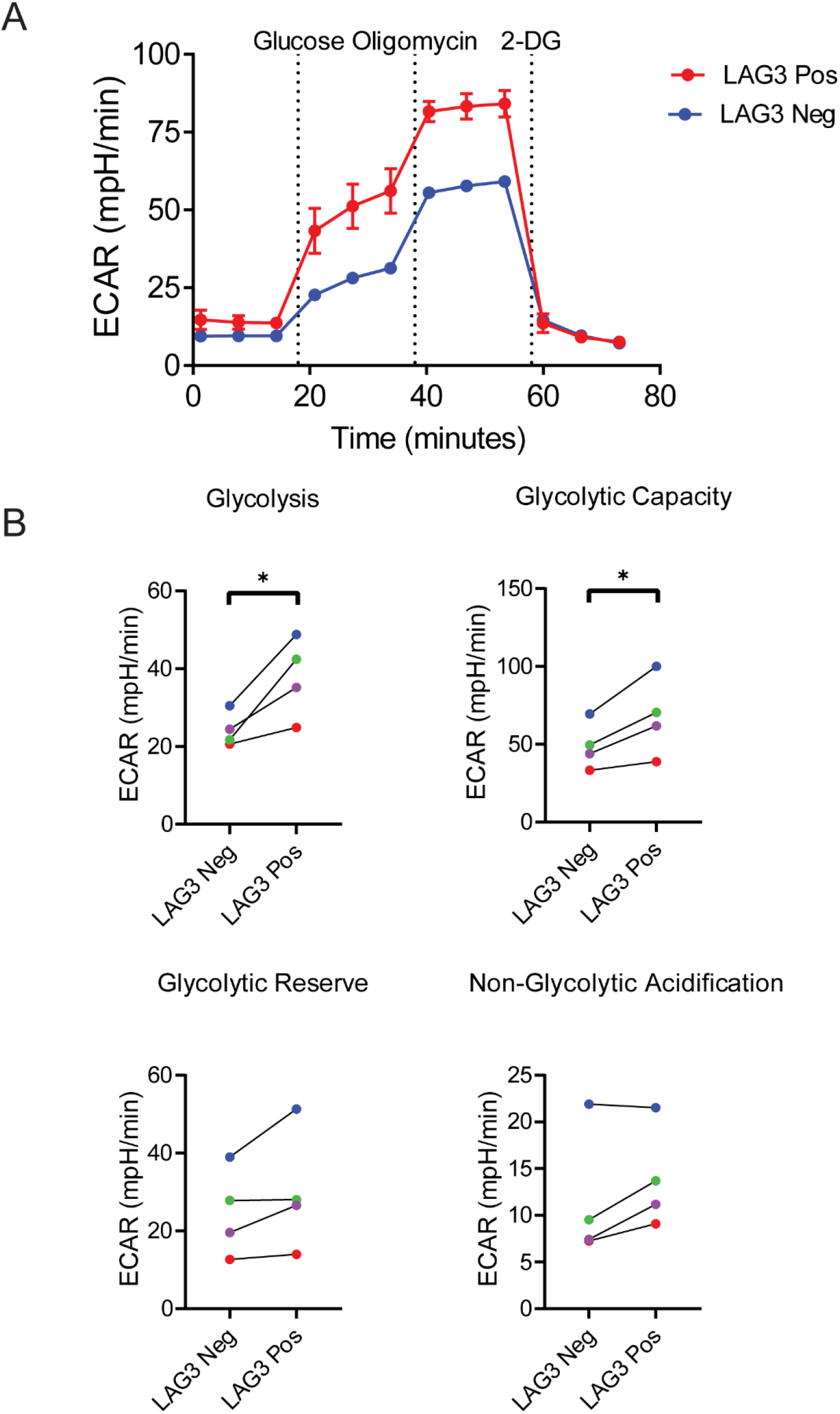
LAG3+ NK cells have higher glycolysis than LAG3-NK cells. **(A)** Representative glycolytic function by ECAR (mpH/min) in in vitro expanded LAG3 positive (red) and negative (blue) cells obtained by sequential addition of glucose, oligomycin and 2-deoxy-glucose (2-DG). **(B)** Graphs show significantly higher glycolysis (left) and glycolytic capacity (right) in LAG3 positive cells. N=4 different donors. Statistical differences were analysed using t-test.

### LAG3 is a negative regulator of NK cell cytokine production

An earlier study found that mice lacking LAG3 in NK cells were incapable of killing target cells, suggesting a critical role for LAG3 in cytotoxic function (10). However, the function of LAG3 in human NK cells remains unclear (11). Therefore, to assess its role in cytotoxicity, NK cells were isolated from PBMCs, primed with IFN-α / IL-2 and then expanded *in vitro* for 3 weeks. These NK cells were cultured with target cells loaded with Calcein-AM dye at different effector to target ratios for 5 hours in the presence of a LAG3 blocking antibody, or isotype control, and the cytotoxicity was measured by the fluorescence of Calcein dye released into the medium upon target cell lysis. Blocking LAG3 did not have any effect on the killing activity of NK cells (Fig 5a). However analysis of cytokine secretion revealed an increase in the production of IFNγ, TNFα, MIP1α, MIP1β and GM-CSF by NK cells when they were cultured in the presence of anti-LAG3 blocking antibody compared to control IgG (Fig 5b). These results demonstrate that LAG3 has a negative regulatory effect in NK cell production of cytokines without significantly affecting their cytotoxic function.

**Figure 5:**
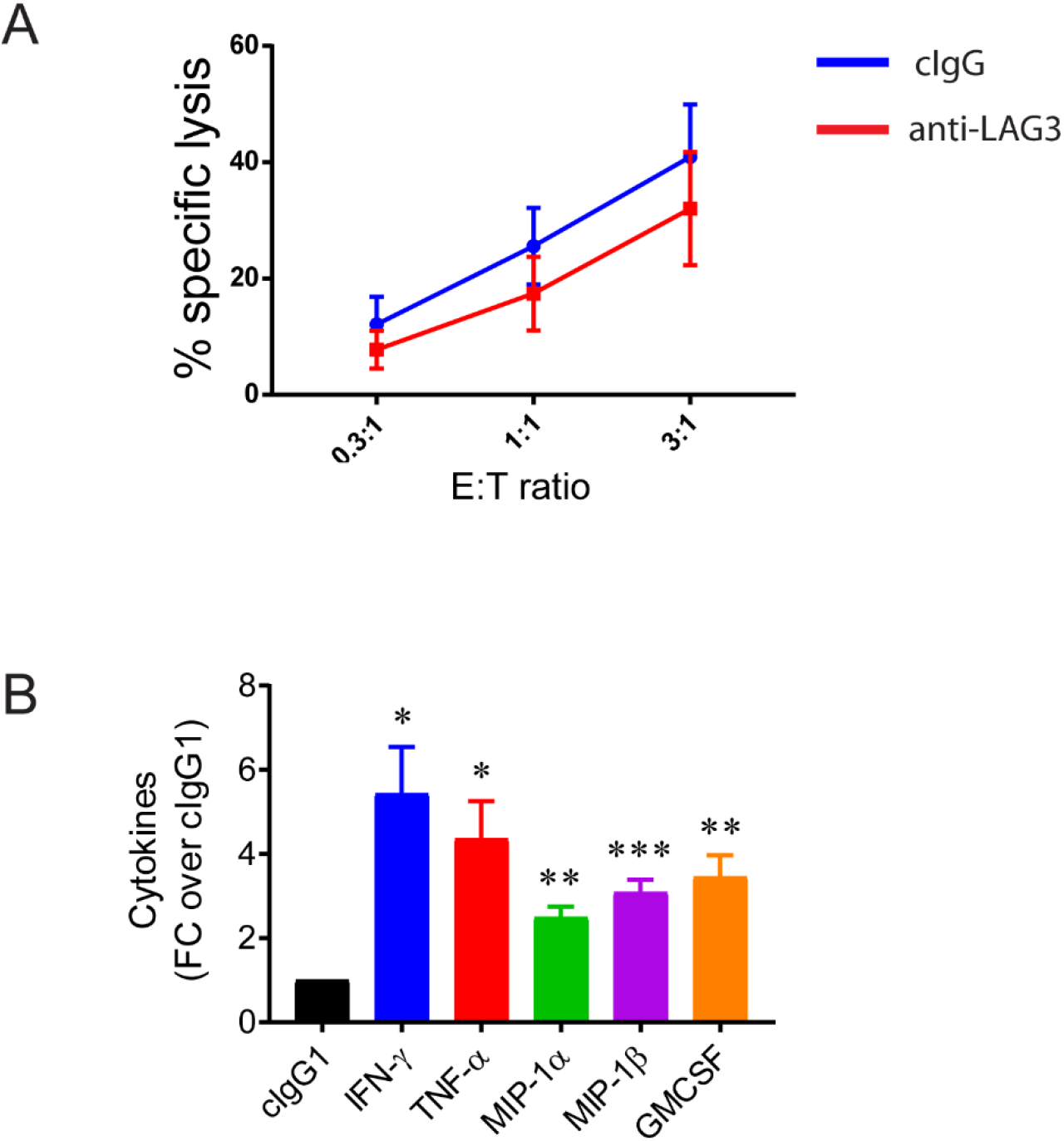
Blocking of LAG3 results in enhanced cytokine production by NK cells. **(A)** NK cells expanded in vitro were co-cultured with target cells loaded with calcein dye at different E:T ratios for 5 hours in the presence of anti-LAG3 antibody or control IgG and analysed for NK cell cytotoxicity. **(B)** The supernatant from the assay were analysed for cytokine secretion by NK cells. The values depicted are fold change of anti-LAG3 over cIgG for each cytokine. Statistical analysis was done using one-way anova with multiple test corrections. N=4 different experiments.

## Discussion

In this study we have established that LAG3 plays a central role in regulating NK cell cytokine production. We also showed that IFN-α rapidly induces the expression of the LAG3 inhibitory receptor on NK cells. Using high dimensional flow cytometry, we characterised LAG3+ NK cells to be both mature and activated based on their expression of CD57, NKG2C and TIM3. Furthermore, LAG3 expressing NK cells also had enhanced glycolytic capacity suggesting a metabolically mature phenotype. Finally, we demonstrated that LAG3 is unique in its ability in selectively regulating inflammatory cytokine secretion and not cytotoxicity. Taken together, these results conclusively characterise the phenotype and function of LAG3 in activated human NK cells.

PEG-IFN-α is an important therapeutic option for a number of infectious diseases and cancer. IFN-α mediates its effects in part through the induction of hundreds of genes (ISGs) which impart anti-viral functions directly, by inhibiting viral replication, and indirectly, through activating innate immune cells. An important cellular effect of IFN-α is the priming of NK cells enhancing their activity. The phenotype and function of the NK cells that are expanded in response to IFN-α treatment is not clear. In this study we showed that exposure to IFN-α induced the expression of LAG3, suggesting that it could play a feedback role in regulating NK cell function. Transcriptomic analyses confirmed an upregulation of LAG3 mRNA by 16 hours. This is in keeping with earlier work demonstrating an increase in LAG3 mRNA and several ISGs in NK cells as early as 6 hours after stimulation with IFN-α suggesting that LAG3 is one of the early expressed genes in response to IFN-α stimulation on NK cells (17).

NK cells are a diverse effector population that vary both in phenotype and function. High dimensional flow cytometry analyses used in our study revealed that LAG3+ NK cells had a higher expression of CD57, a known maturation marker (18). Our analysis also showed that LAG3+ cells expressed increased levels of NKG2C, an activation receptor found to be enhanced in chronic viral infections (18,19). Previous studies have shown that activation, differentiation and function of different immune cells are directed by changes in metabolism (20–23). Metabolic analysis revealed that LAG3+ NK cells have a distinct metabolic profile with enhanced glycolytic reserve and capacity. Interestingly, upon priming with IFNα, LAG3 was predominantly expressed on CD56low NK cells associated with having enhanced cytotoxicity and decreased cytokine secretion compared to CD56bright NK cells. This suggests a mechanism by which the two different arms of NK cell function viz, cytotoxicity and cytokine secretion are regulated independently. It is possible that the expression of several inhibitory receptors like LAG3, KIR and TIM3 assist in keeping effector NK cells in check. The fact that blockade of LAG3 resulted in enhanced cytokine production supports the idea that its function is to attenuate the cytokine production of mature CD56low NK cells without affecting their killing ability.

Previous work has shown that LAG3 is check point inhibitory receptor expressed on exhausted cytotoxic T cells (24,25). NK cells expressing NKG2C and CD57 were described following chronic stimulation of activating receptors. These exhibited an exhausted phenotype and co-expressed PD-1, TIM3 and LAG3 (26,27). In our analysis a small subset of NK cells expressing PD-1 and TIM3 inhibitory receptors also co-expressed LAG3. But the majority of LAG3 expressing NK cells lacked PD-1, a well-known marker of T and NK cell exhaustion. Only a small proportion of NK cells exhibited exhausted phenotype expressing LAG3, PD-1 and TIM-3. These findings suggest that LAG3 could play other regulatory roles in controlling NK cell function without being directly associated with exhaustion.

An evaluation of LAG3 function in mice revealed a key role in the mediating the cytotoxicity function of NK cells (10). In contrast, the data presented here, and the work of Huard and colleagues(11), show that inhibition of human LAG3 did not affect NK cytotoxic function revealing distinct NK cell functions between humans and mice. Extending this observation, we show here that though LAG3 is preferentially expressed on CD56dim NK cells it acts specifically as a negative regulator of cytokine production. Taken together our results show that IFN-α induced LAG3 acts as a negative regulator of cytokine production without compromising cytotoxic function of NK cells.

## Acknowledgements

The authors would like to acknowledge Anshula Alok and Richard Hopkins for their critical reading of the manuscript and comments. The authors also like to thank Robert Balderas, Keefe Chee, William Wong, and John Wotherspoon from Becton Dickinson and Company for their help with flow cytometry panel design.

## Ethics Statement

The study complied with the Declaration of Helsinki, and the protocol was reviewed and approved by the National Healthcare Group Domain Specific Review Board, Singapore (NHG DSRB Ref: 2000/00828). Written informed consent was acquired from all participants.

## Author Contributions

SN and JC led the design of the study. SN, PA, VB and NK conducted the experiments, analysed the data and drafted the manuscript. SL, AF and JC were involved in data analysis and interpretation. JC, AF, SL and CW participated in the design of the study and helped to draft the manuscript. All authors read and approved the final manuscript.

