## Supplementary data for "LAG3 is a Central Regulator of NK Cell Cytokine Production"

### *Supplementary Material*

#### Supplementary Figures

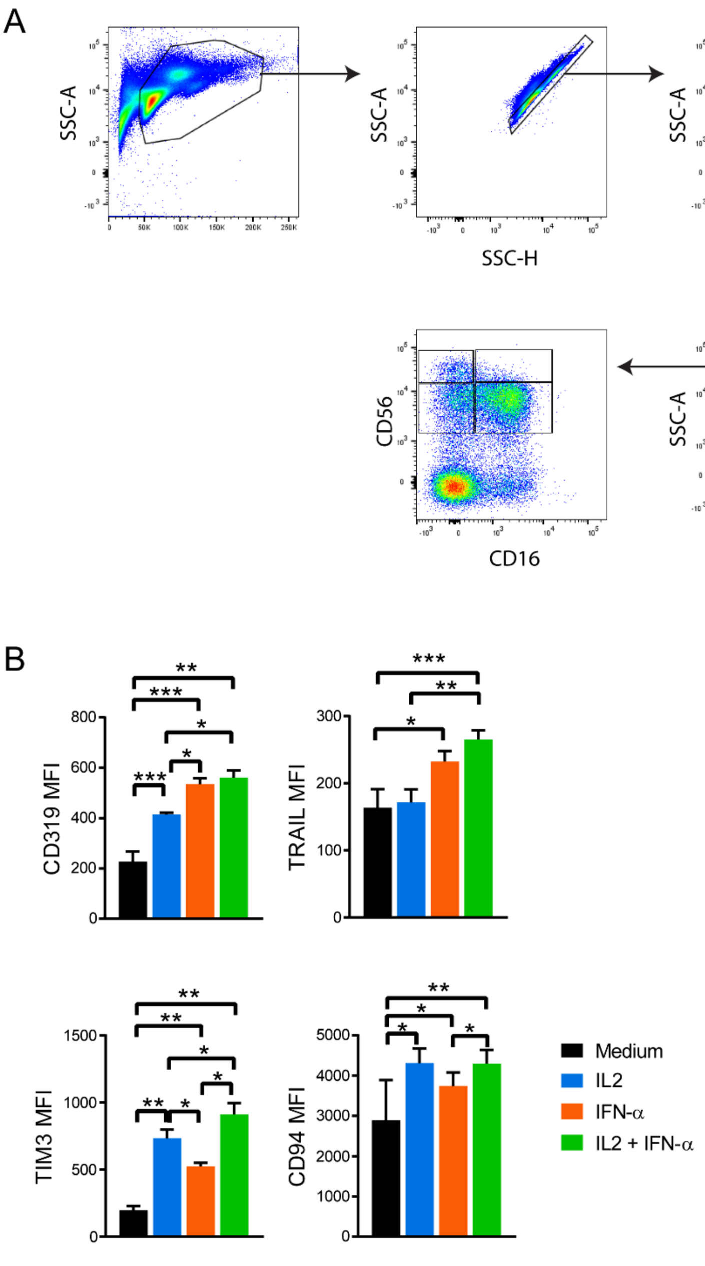


**Supplementary Figure 2.** *Flow cytometry gating for NK cells and NK cell subsets and analysis of NK receptor expression.* **(A)** PBMCs were gated to remove doublets, monocytes, B cells and dead cells. NK cells were then gated based on CD3 and CD56 markers. NK cell subsets were gated using CD16 and CD56 markers. **(B)** Expression of CD319, TRAIL, and CD94 on NK cells stimulated as shown in the legend. N=5 different donors.

A

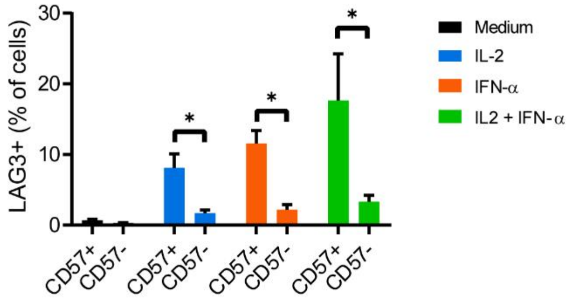


B

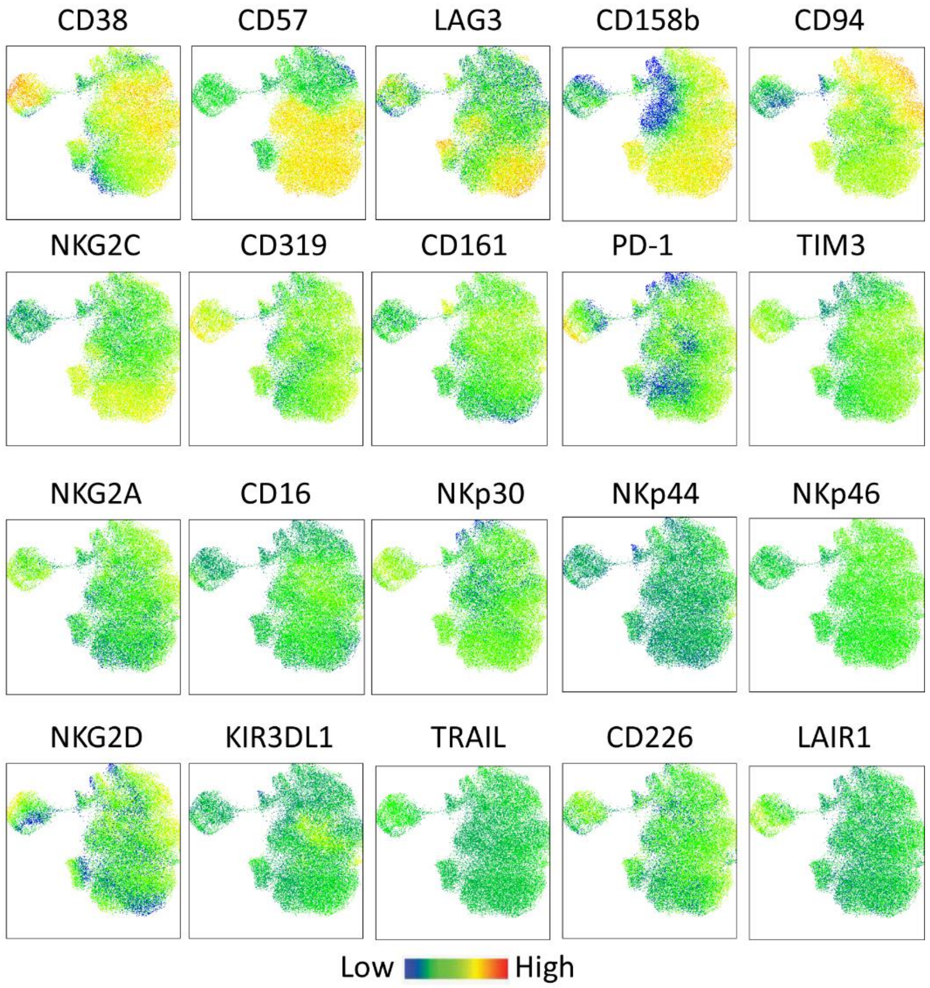


**Supplementary Figure 3.** *Phenotype of LAG3+ NK cells.* **(A)** Frequency of LAG3+ NK cells within CD57+ or CD57- subsets. Statistical difference was analysed by Mann-Whitney test. N=5 different donors. **(B)** UMAP embedding of the clusters coloured according to the levels of expression of various markers from the flow cytometry panel.

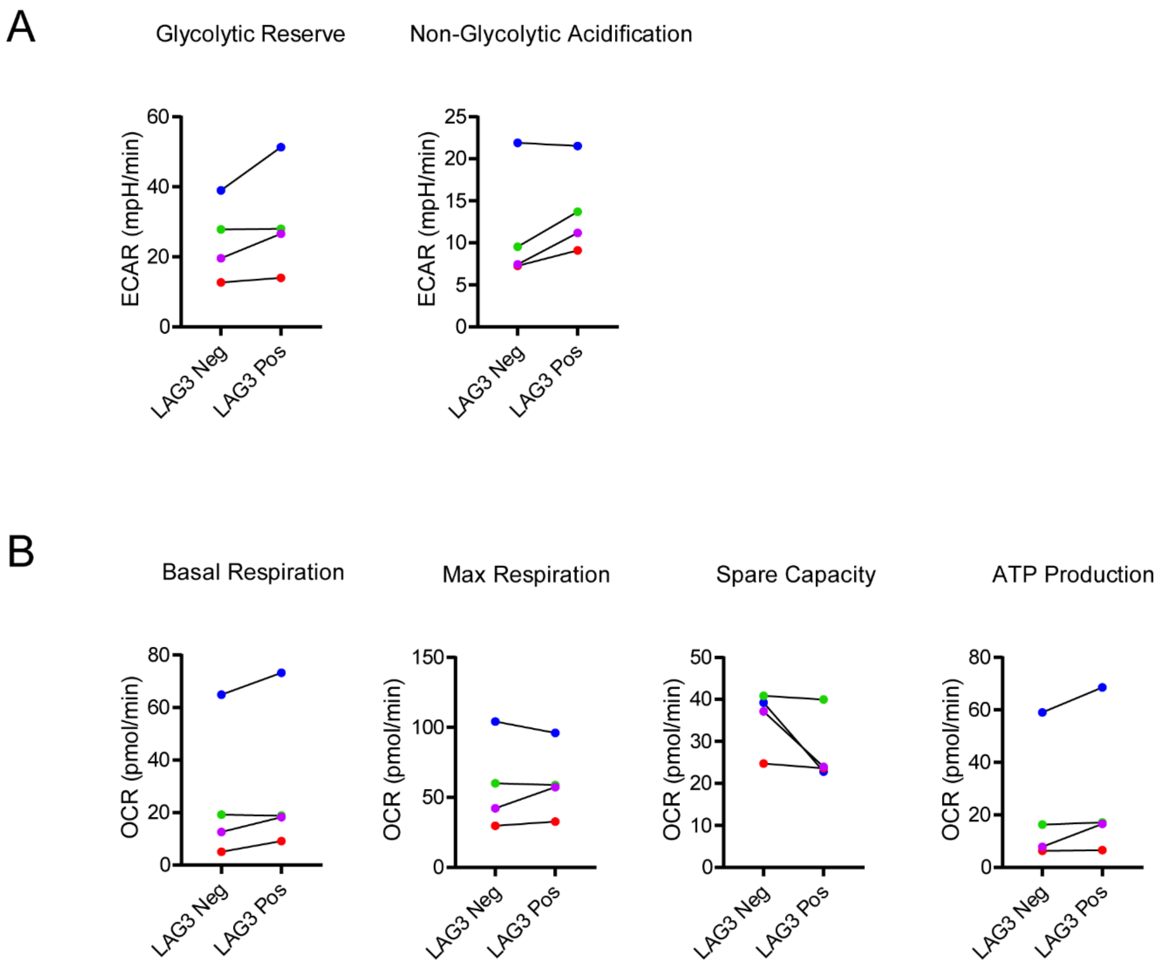


**Supplementary Figure 4.** *Metabolic analysis of LAG3+ and LAG3- NK cells.* **(A)** Characterization of glycolytic reserve (left) and non-glycolytic acidification (right) in LAG3+ and LAG3- NK cells. **(B)** Characterization of mitochondrial oxidative phosphorylation in sorted LAG3+ and LAG3- NK cells. The basal and maximal mitochondrial respiration, spare respiratory capacity and ATP production were measured. N=4 different donors. Statistical differences were analysed using t-test.
